# Tissue-Specific Biochemical Differences Between Chronic Wasting Disease Prions Isolated From Free Ranging, White-Tailed Deer *(Odocoileus virginianus)*

**DOI:** 10.1101/2021.07.19.452961

**Authors:** Kaitlyn Wagner, Robyn Pierce, Elizabeth Gordon, Jennifer R. Ballard, Julie A. Moreno, Mark D. Zabel

## Abstract

Chronic wasting disease (CWD) is an invariably fatal prion disease affecting cervid species world-wide. Prions can manifest as distinct strains that can influence disease pathology and transmission. CWD is profoundly lymphotropic and most infected cervids likely shed peripheral prions replicated in lymphoid organs. However, CWD is a neurodegenerative disease and most research on prion strains has focused on neurogenic prions. Thus, a knowledge gap exists comparing prions in the brain to prions in the lymph node. In this study we compared prions from the obex and lymph node of naturally exposed white-tailed deer to identify potential biochemical strain differences. Here, we report biochemical evidence of strain differences between the brain and lymph node from these animals. Future work should examine the biological and zoonotic impact of these biochemical differences and examine more cervids from multiple locations to see if these differences are conserved across species and locations.

## Introduction

Chronic wasting disease (CWD) is unique among prion disease as the only prion disease known to infect and be naturally transmitted between both captive and free-ranging populations. Mathematical models indicate that direct and indirect horizontal transmission of CWD is the most prevalent form of transmission(1–6), but vertical transmission also contributes to CWD transmission(7–9). Infectious prions have been detected in excreted bodily fluids including saliva, urine and feces, as well as antler velvet, blood and reproductive tissues(7,10– 13). While CWD causes a neurological disease, CWD prions are profoundly lymphotropic and these peripheral prions are the most likely shed into the environment and contribute to horizontal and vertical disease transmission(14–18). Thus, it is critical to determine if any unique characteristics of extraneural prions exist that affect CWD pathogenesis and transmission.

While all prion diseases result from a misfolding of the normal host protein, PrP^C^, to a misfolded form, PrP^Sc^, different biochemical characteristics and disease phenotypes suggest a phenomenon of prion strains transmitting distinct disease characteristics epigenetically enciphered within unique prion structures(19–21). Thus, different prion strains sometimes have significant strain differences. There are multiple CWD strains that have been identified from North American isolates, including CWD-1, CWD-2, H95+ and Wisc-1(22,23). Importantly, the H95+ strain has been shown to emerge from white-tailed deer (*Odocoileus virginianus*) that have the more resistant genotype, 96SS, and this strain has been demonstrated to have an expanded host range, highlighting the importance of continued strain characterization(23). CWD-resistant *PRNP* polymorphisms have emerged in wild elk populations, like Rocky Mountain Elk (*Cervus canadensis nelsoni*), but it is unclear how these polymorphisms may affect novel and/or atypical prion strain emergence in these population(24); as happened with the emergence of the Nor98 scrapie strain in sheep expressing classical scrapie-resistant genotypes(25). Selection of CWD-resistant genotypes in white-tailed deer may also be occurring(26). These data highlight the necessity of continued strain identification and characterization from multiple sources. Understanding strain differences and potentially different transmission dynamics is of critical importance to understand CWD and control its spread.

Of particular concern, extraneural prions have been shown to have increased zoonotic potential(27). This has important implications for cross species transmission and risk for humans potentially contracting CWD from eating infected skeletal muscle or while cleaning a deer in the field (28). While no evidence currently supports natural xenotransmission of CWD prions from cervids to other animal species, CWD prions are infectious to cattle(29), sheep(30), swine(31), and cats(32) when experimentally inoculated intracerebrally. Cattle and cats were resistant to CWD infection after oral exposure, but pigs were susceptible at low levels(31). These data suggest that there is a risk of transmission to additional species and populations, warranting continued monitoring and surveillance of CWD prion strains.

Most of the CWD and prion research completed to date focused on brain-derived prions, likely because prion diseases are neurodegenerative, brain samples are easy to work with and contain the highest titers of prions in infected animals. However, prions shed into the environment likely are extraneural prions, such as those replicated within lymph nodes (LNs). Far less is known about the transmissibility of these peripheral prions, but research suggests LN-derived prions have similar titers to brain-derived prion titers on transgenic mouse bioassay (33). Furthermore, numerous immune receptors and different proteins involved in the complement cascade have been shown to influence prion strain selection, implicating the immune system as an important player in prion strain selection (34–40). This research suggests that lymphogenic prions likely exhibit more strain diversity than neurogenic prions (41,42). Tissue-specific differences in strain heterogeneity, as reflected in the prion cloud hypothesis, predict different prion strains with different biochemical and structural characteristics, in LNs than in the brain(41). Therefore, any differences between the brain-derived and lymph node-derived prions must be investigated to aid our understanding of intra-host and inter-species prion dynamics.

Based on current knowledge of CWD transmission, prion strain selection and differential interspecies transmission, we hypothesize that LNs replicate more diverse CWD prion strains than the brain within and among individuals. While extensive research has focused on brain samples from cervid and transgenic mouse brains, less research has been dedicated to studying and characterizing peripheral prions, leaving a critical knowledge gap that this work addresses. Furthermore, very little work has characterized structural differences between brain and lymph node derived prions from a natural host prior to passage to transgenic mice or other model organisms. While bioassay is a central pillar to prion biology and strain characterization, there are other host factors and transgene expression level differences that influence strain emergence, emphasizing the importance of assessing strain characteristics of prions isolated from the natural host(43,44).

For this study, we assessed biochemical strain differences between paired obex and lymph node samples from naturally exposed white-tailed deer from Arkansas, USA. These analyses reveal significant differences between brain-derived and lymph node-derived prion isolates in some of our biochemical assays. While we observed no conformational stability differences between brain- and the lymph node-derived prions, we observed electrophoretic differences and statistically significant differences in the glycoform ratio of PrP^Sc^ from brain compared to lymph-node samples. Lymphogenic prions exhibited greater overall variance in mean glycoform ratios and conformational stability than neurogenic prions. Surprisingly, we observed greater biochemical differences among brain-derived prions than LN-derived prions across individuals. These data lead us to propose a mechanism whereby the lymphoreticular system propagates a diverse array of prions from which the brain selects a more restricted pool of prions that may be quite different than those from another individual of the same species. Assessing differences in biochemical signatures between prions from brain and lymph nodes among individuals will inform future studies poised to assess biological differences, including zoonotic potential, between, neurogenic and lymphogenic prions using bioassay and other traditional prion assays.

## Materials and Methods

### Sample homogenization

Lymph node and obex samples from white-tailed deer that tested positive by ELISA performed in a National Animal Laboratory Network lab for chronic wasting disease (CWD) were provided frozen from the Arkansas Game and Fish Commission. Samples were stored at -20°C until processing. We implemented several measures to minimize sample cross-contamination. Samples were trimmed with disposable scalpel blades on a half of a petri dish. Both were discarded after a single use. Gloves and lab bench paper were also changed between each sample. Lymph node samples were then placed in homogenizing tubes (product number) with 7-10 zinc zirconium homogenizing beads (2.3 mm diameter) and homogenized to 20% w/v in PMCA I buffer (1x PBS 150 mM NaCl, 4 mM EDTA) with cOmplete protease inhibitor (Roche). Samples were homogenized on a BeadBlaster for 10 rounds, with each round consisting of 3 cycles of a 30 sec pulse at 6 m/s followed by a 10 sec rest between each pulse. Samples were rested on ice for 5 min between each of the 10 rounds. Once samples were homogenized, samples were aliquoted and stored at -20°C until further use. Obex samples were also processed to 20% w/v homogenate in PMCA I buffer and protease inhibitor as described above, but obex samples were homogenized with 7-10 glass beads (2.7 mm diameter) and for 2-3 rounds with a 5 min rest on ice between each round on the BeadBlaster. Samples were then aliquoted and stored at -20°C until use.

### Conformational Stability Assay (CSA) and Glycoform Ratio

To assess the conformational stability of the prions from the brain and the lymph node, samples were thawed and 15 µl of sample was added to 15 µl of GndHCl in 0.5 M increments from 0-4 M, briefly vortexed and incubated at room temperature for 1 hour. After the 1 h denaturation, samples were precipitated in ice-cold methanol overnight at -20°C. The following day, samples were removed from the -20°C, centrifuged at 13,000 rcf for 30 min at 4°C. Then, GndHCl and methanol were removed, and the protein pellet was resuspended in either 18 µl of PMCA I buffer (lymph node samples) or 36 µl of PMCA conversion buffer (1x PBS 150 mM NaCl, 4 mM EDTA, 1% Triton-X 100, obex samples). Lymph node samples then had 2 µl of 500 µg/mL of proteinase K (PK, Roche) (diluted in 1x PBS and 0.5 M EDTA) added for a final PK concentration of 50 µg/mL in each sample. Obex samples had 4 µl of 1000 µg/mL of proteinase K (PK, Roche) (diluted in 1x PBS and 0.5 M EDTA) added for a final PK concentration of 100 µg/mL. Samples were then incubated on a shaking heat block for 30 min @ 37°C and 800 rpm. Twenty microliters of each sample were denatured in the presence of 10 µl of 3x loading buffer (2.5 volumes of 4x sample loading buffer [Invitrogen] per 1 volume of 10x sample reducing agent (Invitrogen) for 10 min at 95°C. Samples were then either saved at -20°C or immediately run by western blot and analyzed for conformational stability and glycoform ratio.

### Western blotting

Samples were run on 12% bis-tris gels [NuPage] in 1x MOPS running buffer and transferred to polyvinylidene difluoride membranes. Non-specific binding was reduced by blocking the membranes in 5% nonfat dry milk and 1% tween-20 in 1x PBS (NFDM) for 1 hour with rocking at room temperature. Membranes were then incubated in HRP-conjugated anti-PrP monoclonal antibody Bar224 (Cayman Chemical) diluted to 1:20,000 in SuperBlock (Thermo Fischer) overnight at 4°C. Blots were washed the following day in PBST (0.2% Tween20 in 1x PBS) six times for 5 minutes each wash. Membranes were developed using enhanced chemiluminescent substrate (Millipore) for 5 minutes before imaging on ImageQuant LAS 4000 (GE).

### Data analysis

Densitometric analyses were completed in ImageJ. Statistical analysis and graphing were performed in GraphPad Prism (version 8.30). Conformational stability was determined by calculating the concentration at which the signal was half of the input ([GndHCl]_1/2_) after fitting the data to a fourth order polynomial regression in GraphPad Prism. Glycoform ratio was calculated in ImageJ by determining what percentage of the total signal was contributed by each glycosylation state. Glycoform ratio data was arcsine transformed before statistical analysis so percent data would fit a normal distribution. Only samples that had at least three successful replicates (conformational stability) or had results replicated on at least two blots with three samples each (glycoform ratio) were included for analysis.

## Results

### Sample Origin, Preparation and Result Overview

Samples used in this study were all collected from naturally exposed white-tailed deer in the state of Arkansas and shared with us from our collaborators from the Arkansas Fish and Game Commission. We optimized our assays to obtain the clearest, most reproducible data possible. If we employed the same PK digestion and Western Blot methods for brain/obex samples and lymph node samples, Brain-derived PrP^Sc^ signals were indistinguishable from PrP^C^ signals. We therefore optimized digestion conditions for each tissue type. Brain samples required PK digestion in the presence of 1% Triton-X 100, with a higher concentration of proteinase K (PK, 100 µg/mL), and less starting total protein (5% w/v homogenate before PK digestion) electrophoresed through the gel (data not shown). Lymph node samples ran well when more protein was loaded onto the get (10% w/v starting homogenate) and digested with less PK (50 µg/mL) than obex samples. Of the nine animals that had paired obex and lymph node samples, only four of the animals gave us interpretable data from both the tissues that enabled us to compare intra-host variation (Table 1).

**Table 1.** Animal identification number and summary of biochemical strain typing results.

| Sample | Age (y) | Sex | Obex |  | Lymph Node |  |
| --- | --- | --- | --- | --- | --- | --- |
|  |  |  | CSA | Glycoform Ratio | CSA | Glycoform Ratio |
| 10023 | 2.5 | F | Yes | Yes | Yes | Yes |
| 10074 | 5.5+ | F | Yes | Yes | Yes | Yes |
| 10083 | 1.5 | M | Yes | Yes | Yes | Yes |
| 07399 | 5.5+ | F | Yes | Yes | Yes | Yes |
| 07416 | 3.5 | M | No | No | Yes | Yes |
| 10030 | 3.5 | M | No | No | Yes | Yes |
| 14707 | 4.5 | M | Yes | Yes | No* | No* |
| 10080 | 2.5 | M | No | No | No | No |
| 07415 | 2.5 | M | No | No | No | No |

*\*only 1 western blot of these samples (of 5 attempts) gave interpretable data. Too few replicates for conformational stability analysis or glycoform ratio to be determined. Samples were excluded from analysis. CSA, conformational stability assay.*

### Conformational differences between obex and lymph node samples at ≥ 2.5 M GndHCl

Samples were prepared for analysis by conformational stability and glycoform ratio as described in the methods section. Samples were then run on a Western blot to collect densitometric and electrophoretic mobility data. Differences in electrophoretic mobility of a prion sample reveals structural differences that dictate PK accessibility, resulting in different PK-resistant core fragments of PrP^Sc^. These heritable structural differences are reliable biochemical indicators of different prion strains (45,46). Obex samples that were incubated in 2.5 M GndHCl and greater migrated faster than samples exposed to lower concentrations of GndHCl (Figure 1). This was only observed in obex samples and these data were consistent among all four individuals.

**Figure 1.**
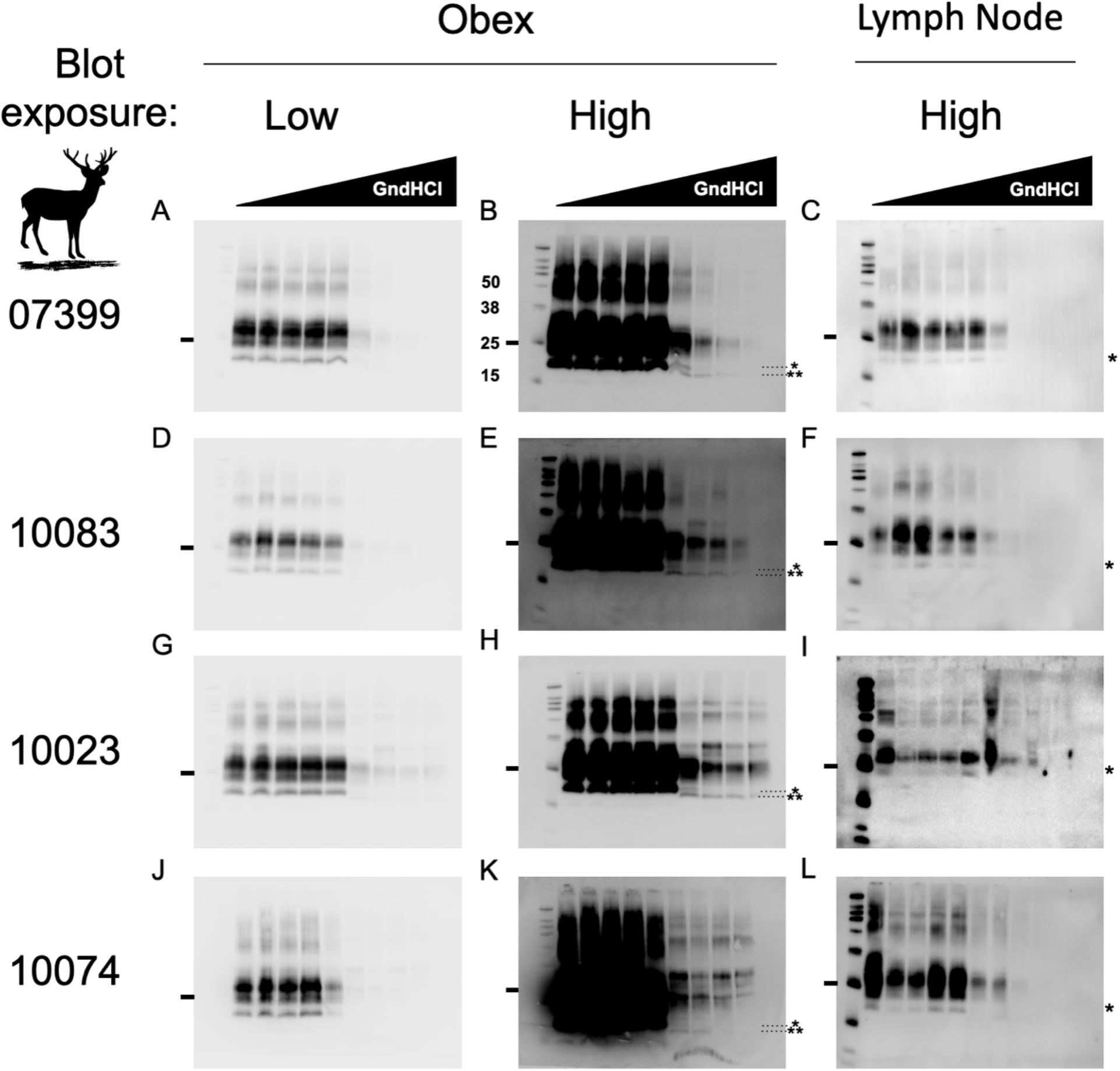
Obex prions adopt an alternative conformation in the presence of ≥ 2.5M GdnHCl compared to lymph node prions from the same animal. Western blots to the right of the animal identification number are all from the same individual. Sample obex blots imaged at a typical exposure (panels A,D,G,J) and overexposed (B,E,H,K) reveal a unique, faster-migrating PrP^Sc^ electrophoretic signature in obex samples (note the unglycosylated 19kD band* at < 2.5 M GndHCl compared to a 17 kD band** at ≥ 2.5M GdnHCl) absent in PrP^Sc^ from lymph node samples (C,F,I,L). Markers to the right of blots indicates the 25 kD molecular weight (MW) band. Other bands from the protein MW ladder, from top to bottom, indicate the 260, 125, 90, 70, 50, 38, 25, 15 and 8 kD (visible only in lymph node blots in panels C,F,I,L) MW markers.

### Greater variability in conformational stability of lymph node prions compared to brain prions

Distinct prion strains can have different conformationally stability in the presence of chaotropic denaturing agents like GndHCl. We compared conformational stability of prions isolated from lymph node to prions isolated from brain samples to determine if this strain characteristic would reveal potentially different strains from different tissues within the same animal. We treated samples with increasing concentrations of GndHCl and determined their [GdnHCl]_1/2_ values, which is the [GdnHCl] that eliminates half of the PrP^Sc^ compared to the untreated sample. While one animal trended toward statistical significance, we detected no statistical differences in conformational stability measured between paired obex-derived and lymph-node derived prions in any of the four individuals examined (unpaired t-test, p<0.05, Figure 2). However, we did observe a statistical difference in the variance of mean [GdnHCl]_1/2_ values between the obex and lymph node-derived prion samples from animal 10083 (F-test, p<0.05). Differences in mean [GdnHCl]_1/2_ variances for animal 10074 trended towards significance (F-test, p=0.07). The other two samples did not exhibit significant difference in conformational stability variance (F-test, p>0.05).

**Figure 2.**
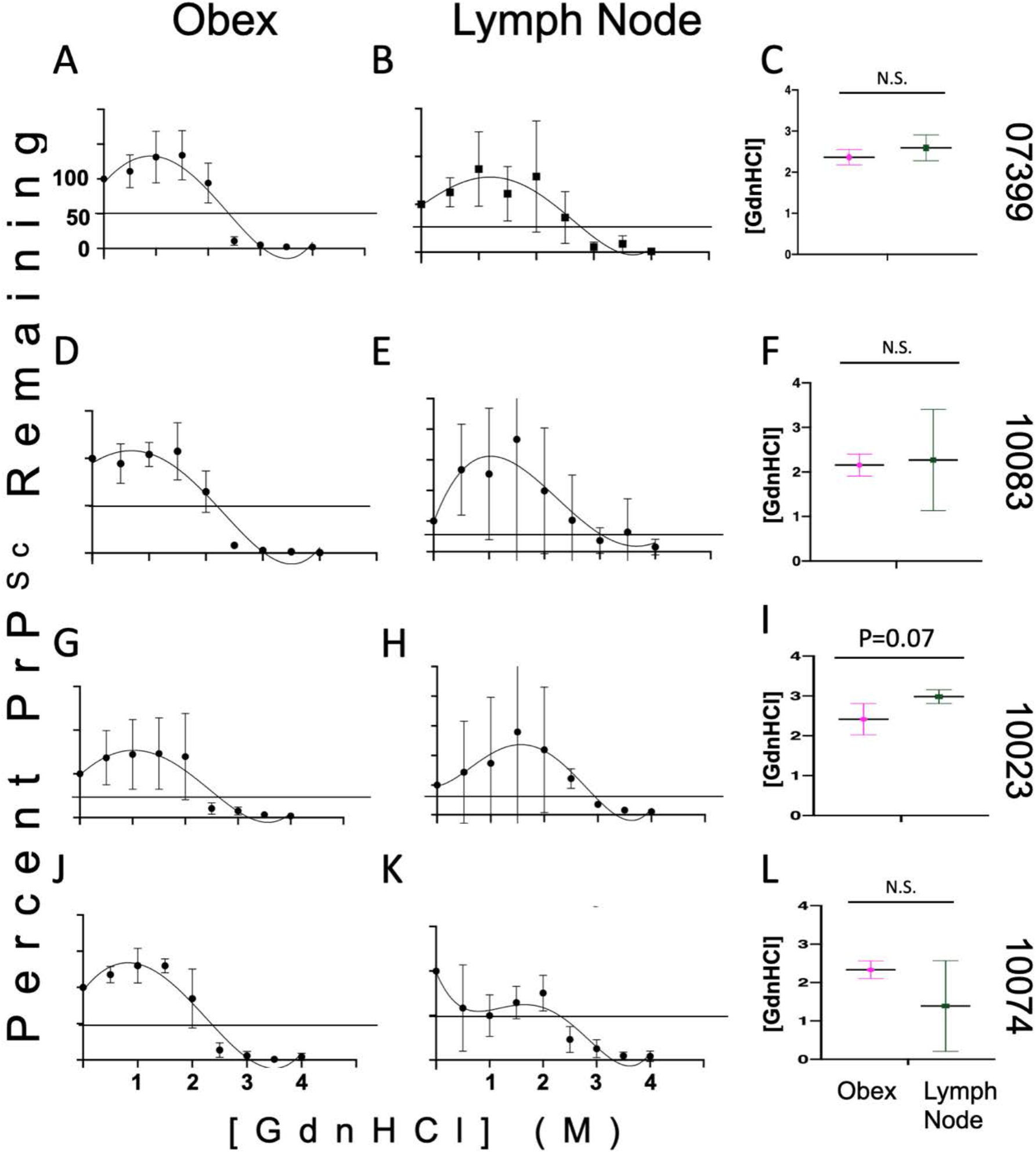
No differences in conformational stability between obex- and lymph node-derived prions in paired samples from the same deer. No differences were observed between obex and lymph node samples from deer 07399 (A, B, C), 10083 (D, E, F), 10023 (G, H, I) or 10074 (J, K, L). Samples were treated with GndHCl as described in the methods section. Fourth-order polynomial regression curves are shown for both obex (A, D, G, J) and lymph node (B, E, H, K) samples and depict the average curve from at least 3 experiments. The curves show the mean and 95% Confidence Interval (CI) of the samples at each concentration of GndHCl. Different y-axes are shown so the data is easily visualized and can more accurately depict the range of the standard deviation for the samples at each concentration of GndHCl. The horizontal line in each graph depicts 50% of the signal remaining compared to untreated samples, set at 100%. Panels C,F,I and L depict the mean and standard deviation of the GndHCl_1/2_ values from the individual replicates for both the obex and the lymph node of the same animal. While no statistical differences were found between sample means, the difference between obex and lymph node of sample 10023 is trending towards significance (p=0.07). Unpaired t-test, p<0.05. N.S., not significant.

To assess potential differences in conformational stability among individuals in either the brain or the lymph node, we compared prions isolated from the same tissue across all individuals that yielded interpretable data, not just the four samples with paired obex and lymph node data that allowed for within animal comparison (Table 1). We observed no statistical differences in conformational stability (mean [GndHCl]_1/2_) in prions derived from either brain or lymph node when compared among individuals (Figure 3). To determine if significant biochemical differences exist between brain and lymph node prions generally across individuals, we analyzed mean [GdnHCl]_1/2_ values calculated from individual values aggregated for each tissue from all individuals. While we observed no significant differences in mean [GndHCl]_1/2_ values from brain (1.9 M, 95% CI: 1.8 - 2M) and lymph node (2.2M, 95% CI: 1.7 – 2.7M; paired t-test, p<0.05), lymph node prion samples exhibited statistically more variance in mean [GndHCl]_1/2_ than obex samples (Figure 4, F-test, p<0.05).

**Figure 3.**
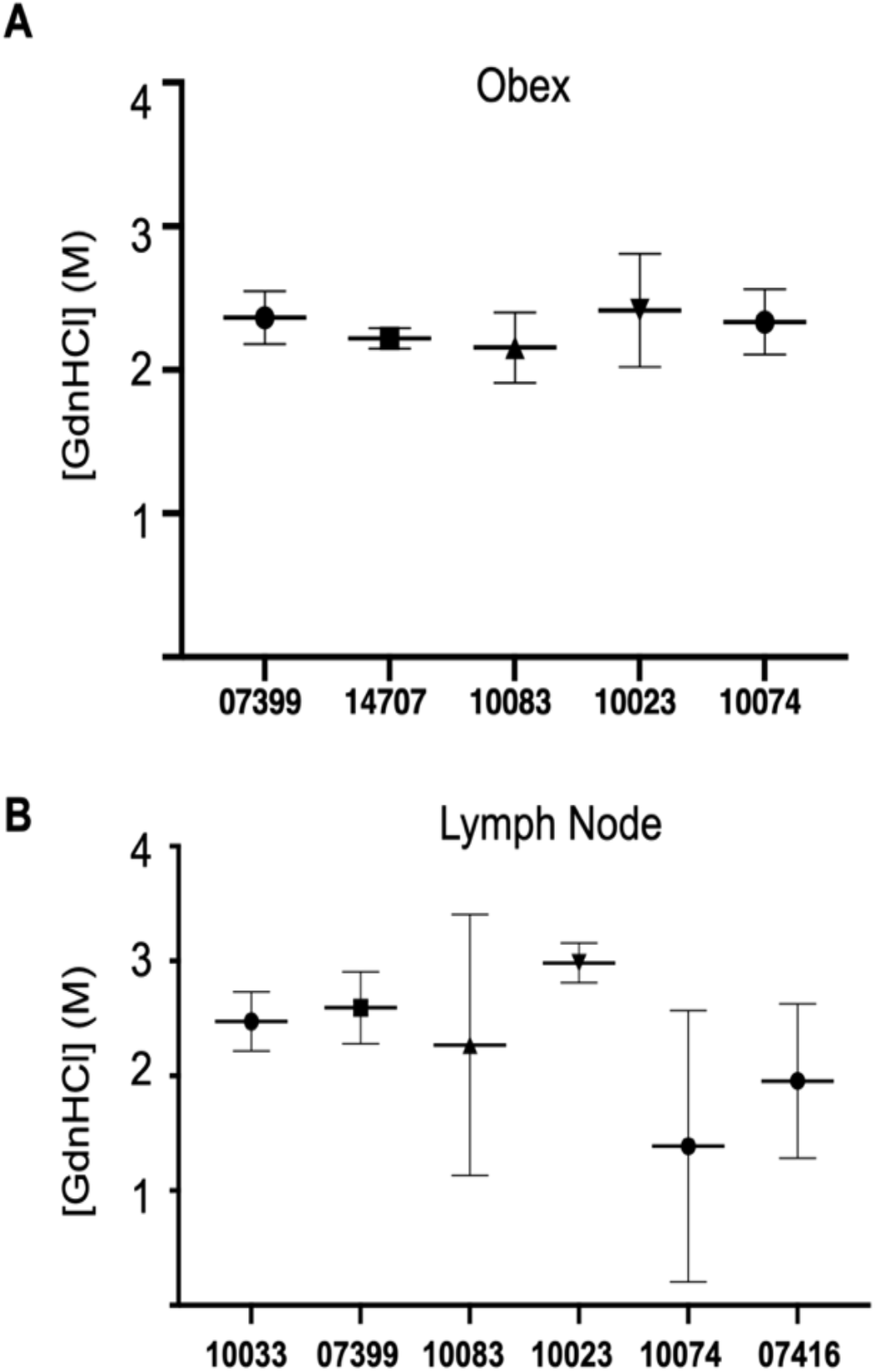
No conformational stability differences among individuals in either obex or lymph node derived prions. We observed no differences in mean [GdnHCl]_1/2_ values in prions isolated from (A) obex samples or (B) lymph nodes among any individuals. One-way ANOVA with Tukey adjustment (p>0.05).

**Figure 4.**
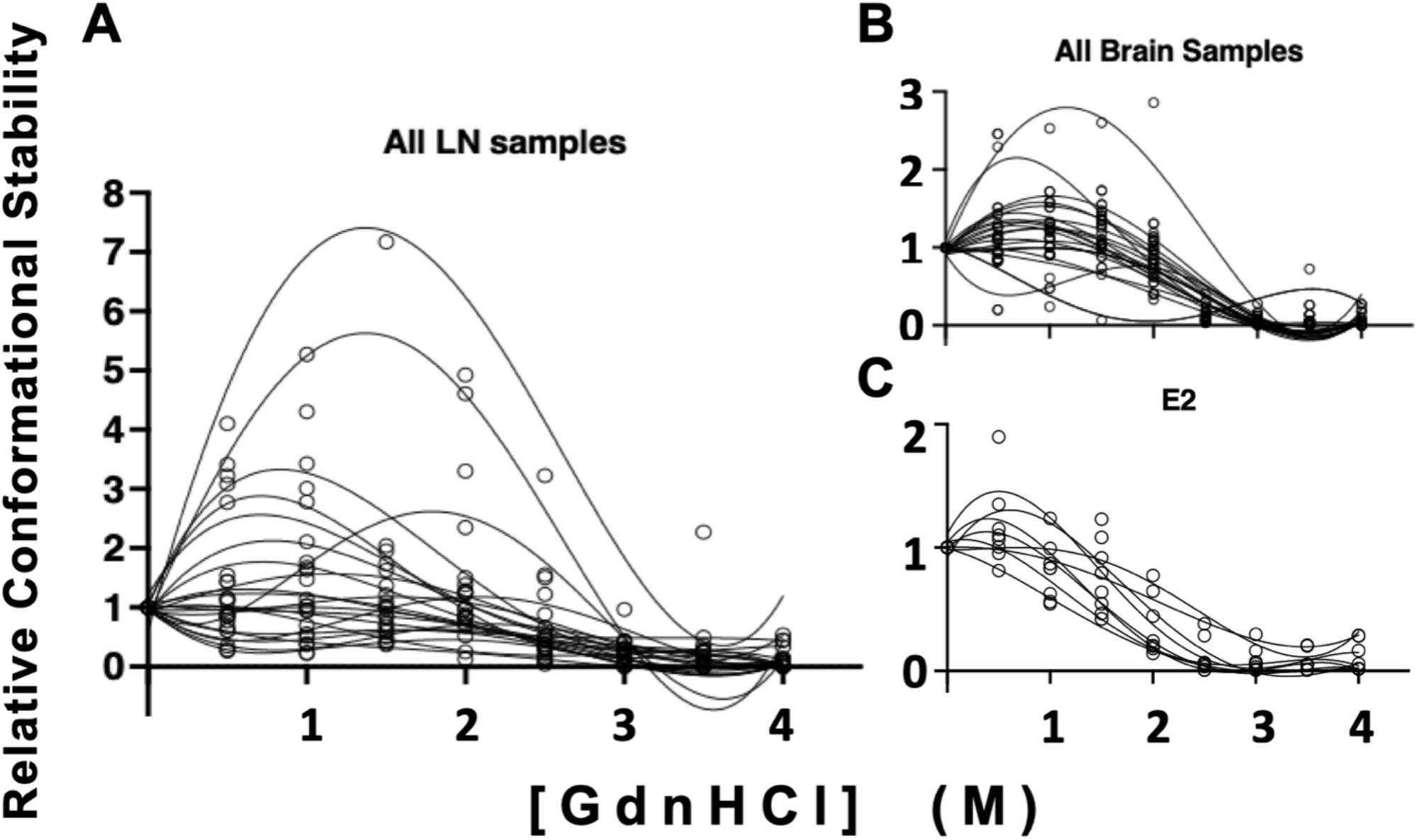
Significant difference in variance of [GdnHCl]_1/2_ values between LN and brain-derived prion isolates across individuals. While we observed no statistical differences in mean [GdnHCl]_1/2_ values between LN and brain derived prion isolates (paired t-test, p>0.05), we found significant differences in the variance of these means (F test, p<0.05). Conformational stability curves for each isolate is shown in triplicate. (A) Prions isolated from LNs exhibited increased variance compared to (B) prions isolated from brains and (C) an Elk brain used as a laboratory control (E2).

### PrP^Sc^ Glycoform ratio differences between prions from paired brain and lymph nodes in the same animal

Glycoform ratios are another heritable biochemical trait of prion strains and have been used in the characterization of the prion strains causing BSE, CJD and CWD (47–49). Just as we assessed conformational stability, we compared PrP^Sc^ glycoform ratios of prions in obex and lymph node tissue samples from the same deer to assess whether distinct prions may reside in distinct tissues within the same host. We found significant differences in proportions of at least two glycoforms between matched brain and lymph node samples for all four individuals examined (unpaired t-test, p<0.05, Figure 5). When comparing glycoform ratios of prions in tissues across individuals, we observed few differences in glycoform ratio in lymph node prions (Figure 5 and Table 2). However, we detected many glycoform ratio differences among obex prions across individuals. In fact, only two obex samples that, when we compared their PrP^Sc^ glycoform ratios to each other, were not statistically different (ANOVA with Tukey adjustment; Figure 5 and Table 3). Finally, when comparing mean glycoform ratios calculated from aggregated individual ratios for each tissue across all individual deer, we observed statistical differences in glycoform ratio across all three glycosylation states (paired t-test, p<0.001; Figure 5C).

**Figure 5.**
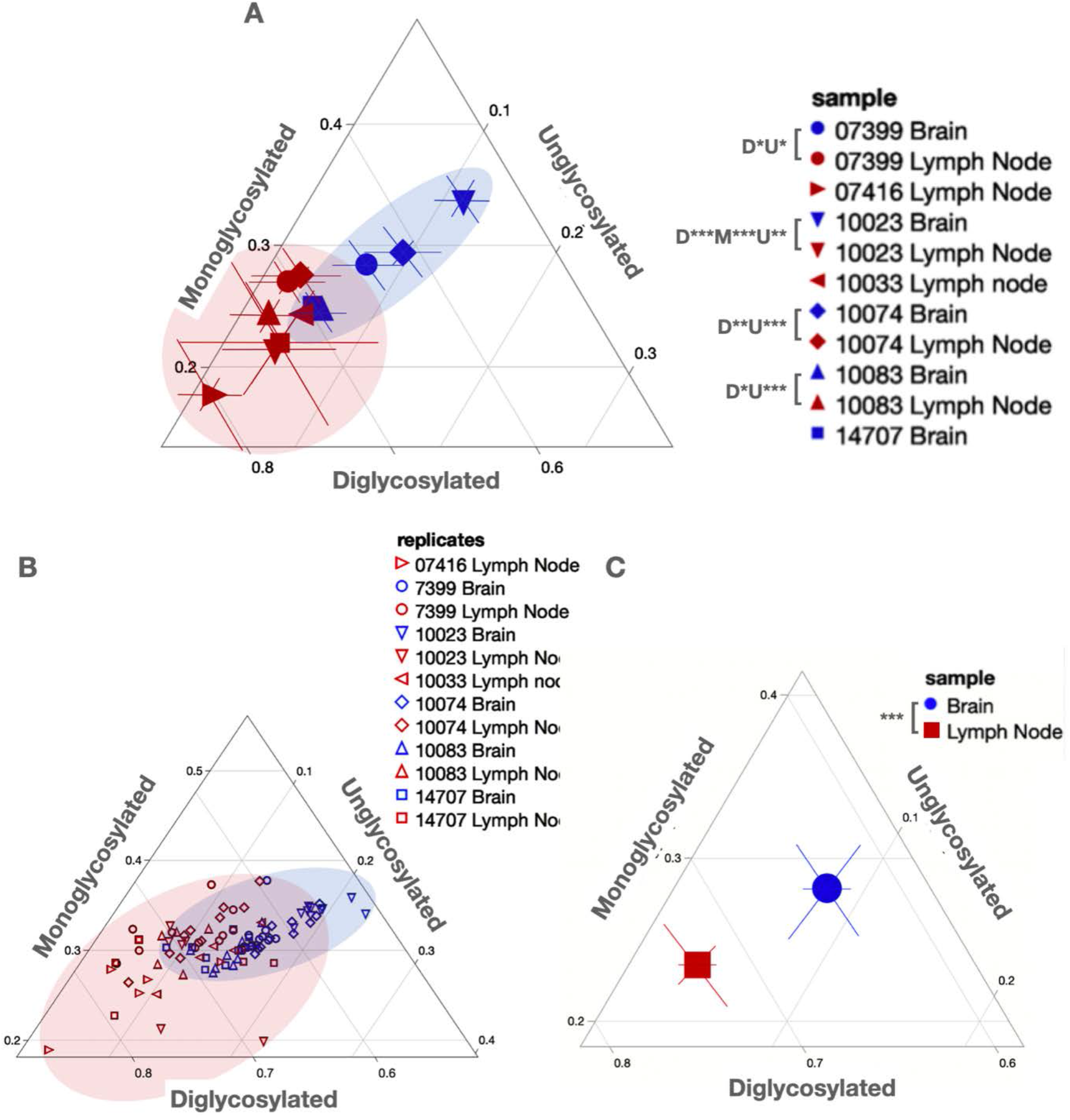
PrP^Sc^ Glycoform ratio differences within and among deer tissue samples. Ternary plots facilitate glycoform ratio comparisons. (A) Mean glycoform ratios with 95% CI shown for paired (0733 (circles), 10023 (down triangles), 10074 (diamonds) and 10083 (up triangles)) and unpaired samples from obex (Brain, blue) and lymph node (red). P-values are shown in the legend for di-(D), mono-(M) and unglycosylated proportions. (B) raw glycoform ratio data shown for all replicates from all tissues analyzed. (C) Overall mean glycoform ratios with 95% CI for all brain (blue circle) and lymph node (red square) samples aggregated from individual glycoform ratios from all replicates from all animals shown in (B). Shaded areas depict the range captured for PrP^Sc^ glycoform ratios from both tissues. *p<0.05, **p< 0.01, ***p<0.001. One-Way ANOVA with Tukey adjustment.

**Table 2.** PrP^Sc^ glycoform comparison in prions across all paired lymph node samples.

| **Samples compared<sup>^</sup>** | Diglycosylated |  | Monoglycosylated |  | Unglycosylated |  |
| --- | --- | --- | --- | --- | --- | --- |
|  | p value | Significance | p value | Significance | p value | Significance |
| 07399 vs. 10083 | 0.9514 | ns | 0.9369 | ns | 0.9996 | ns |
| 07399 vs. 10023 | 0.7472 | ns | 0.1899 | ns | 0.5652 | ns |
| 07399 vs. 07416 | 0.0124 | * | 0.0107 | * | 0.9990 | ns |
| 07399 vs. 10033 | >0.9999 | ns | 0.9204 | ns | 0.2456 | ns |
| 07399 vs. 10074 | 0.9979 | ns | >0.9999 | ns | 0.8841 | ns |
| 10083 vs. 10023 | 0.9991 | ns | 0.8572 | ns | 0.8754 | ns |
| 10083 vs. 07416 | 0.2354 | ns | 0.2354 | ns | 0.9926 | ns |
| 10083 vs. 10033 | 0.9900 | ns | >0.9999 | ns | 0.5846 | ns |
| 10083 vs. 10074 | 0.8418 | ns | 0.8949 | ns | 0.9932 | ns |
| 10023 vs. 07416 | 0.3617 | ns | 0.8279 | ns | 0.5411 | ns |
| 10023 vs. 10033 | 0.9168 | ns | 0.8781 | ns | 0.9907 | ns |
| 10023 vs. 10074 | 0.5528 | ns | 0.1598 | ns | 0.9798 | ns |
| 07416 vs. 10033 | 0.0687 | ns | 0.2556 | ns | 0.2635 | ns |
| 07416 vs. 10074 | 0.0065 | * | 0.0092 | ** | 0.8289 | ns |
| 10033 vs. 10074 | 0.9964 | ns | 0.8735 | ns | 0.7688 | ns |

<sup>^</sup>One-way ANOVA with Tukey adjustment. \*significant, ns, not significant

**Table 3.** *PrP*^*Sc*^ glycoform comparison in in prions across all paired obex samples.

| **Sample comparison<sup>^</sup>** | Diglycosylated |  | Monoglycosylated |  | Unglycosylated |  |
| --- | --- | --- | --- | --- | --- | --- |
|  | p value | Significance | p value | Significance | p value | Significance |
| 07399 vs. 14707 | 0.0066 | ** | 0.0456 | * | 0.0176 | * |
| 07399 vs. 10083 | 0.0173 | * | 0.0251 | * | 0.3450 | ns |
| 07399 vs. 10074 | 0.3187 | ns | 0.9244 | ns | 0.0678 | ns |
| 07399 vs. 10023 | <0.0001 | **** | 0.0011 | *** | <0.0001 | **** |
| 14707 vs. 10083 | 0.9968 | ns | 0.9993 | ns | 0.6447 | ns |
| 14707 vs. 10074 | <0.0001 | **** | 0.0051 | ** | <0.0001 | **** |
| 14707 vs. 10023 | <0.0001 | **** | <0.0001 | **** | <0.0001 | **** |
| 10083 vs. 10074 | <0.0001 | **** | 0.0025 | ** | 0.0003 | *** |
| 10083 vs. 10023 | <0.0001 | **** | <0.0001 | **** | <0.0001 | **** |
| 10074 vs. 10023 | 0.0022 | ** | 0.0119 | * | 0.1001 | ns |

<sup>^</sup>One-way ANOVA with Tukey adjustment. \* $p < 0.05$ , \*\* $p < 0.01$ , \*\*\* $p < 0.001$ , \*\*\*\* $p < 0.0001$ , ns, not significant

## Discussion

Chronic wasting disease is an invariably fatal disease infecting cervids world-wide. This disease is devastating to the individual animals that become infected and has resulted in substantial population-level effects in free-ranging animals, including population decline and herd culling as a method of disease control(24,50–54). The unique nature of prions and prion diseases, coupled with extremely facile animal-to-animal transmission, necessitates a thorough understanding of the pathogen causing disease.

Because CWD and other prion diseases are neurological, and the nervous system harbors the highest prion titers, most prion research has focused on brain-derived prions. While these pivotal studies are critical to our current understanding of prion disease, the profound lymphotropism and presence of infectious prions in extraneural sites should be considered when investigating environmental CWD prion contamination and indirect CWD transmission. Furthermore, most studies focused on prion strain characterization utilized mouse bioassay, where prion isolates are passaged into transgenic mice expressing PrP^C^ from another species. The resulting disease phenotype in the mouse and the biochemical characteristics of the prions from the brains of infected mice are assessed to give a set of disease characteristics that are then defined as a prion strain(19,20). However, prion strains biologically cloned via mouse passage may be quite different than prions originally isolated from the natural host. Indeed, biological cloning of prions, by definition, results in stabilization of a set of biological and biochemical traits that may be different than those of the original isolates. Biochemical analyses of primary CWD prions isolated from different tissues in the same natural host are essential to understand critical features of prion strains and the diseases they cause. However, scant research so far has investigated thoroughly the biophysical and biochemical characteristics of prion strain isolated from different tissues within and among hosts before passaging these isolates into mice. In most cases, biologically cloning CWD strains by serial passage in the natural host is logistically and financially impossible. Here, we compare biochemical, conformational and stability traits between lymph node and brain-derived prions within and among natural hosts before passaging into mice. Any differences we find may have important implications for horizontal, indirect and even zoonotic CWD transmission and disease progression.

Of the paired samples received for analyses from nine deer, we obtained interpretable data from only four pairs of obex and lymph node tissues for intra-animal comparison (Table 1). We optimized our tissue preparation and Western blotting methods to obtain comparable, interpretable data, highlighting an important note about strain differences observed between the obex and lymph node samples. The central nervous system expresses the most PrP^C^, followed by lymphoid tissues and to a much lesser extent other peripheral tissues (55–63). PK may fail to fully digest high PrP^C^ levels in the brain to give accurate results if not more aggressively digested. Cervid PrP^Sc^ also may aggregate into denser plaques that protect cerPrP^C^, necessitating extra dilution, detergent and/or PK for complete PrP^C^ solubilization and digestion from brain samples.

We had incomplete data for samples from deer 07416 and 10030 because we detected cerPrP^Sc^ only in lymph node, not obex samples. CWD prions often replicate to detectable levels in lymphoid tissues before the brain(14,18,64). So these two brains likely contain too few prions, or perhaps nascent, protease-sensitive oligomeric prions that PK and WB fail to detect. Mouse bioassay could determine the prion titer, if any, and the biological characteristics of these prions, if present, in the brains of these two animals. Surprisingly, we detected cerPrP^Sc^ in obex and lymph node from a male deer aged just 1.5 years (10083). This rare occurrence of prions in the central nervous system (CNS) in this young cervid may indicate an early, perhaps vertical transmission event (7,8,65). We were less surprised to detect cerPrP^Sc^ in lymph node but not brains than our converse discovery. We detected cerPrP^Sc^ consistently in the brain of deer 14707, but inconsistently in the paired lymph node sample. Difficulty with tissue homogenization and western blotting likely contributed to the inconsistent results with that sample. Finally, samples from two deer (10080 and 07415) did not yield interpretable data for either the brain or the lymph node. These samples likely had prion levels that were below the limit of detection by western blot and were not able to be analyzed in this study.

For the four samples that we did have interpretable, reproducible data in both tissues (10023, 10074, 10083 and 07415) we observed no significant differences in the mean [GndHCl]_1/2_ values between the obex- and lymph node-derived prion samples (Figure 2), suggesting similar conformational stability of each isolate. CWD is traditionally one of the more stable prion strains and it may be that a characteristic of CWD overall is that they are very stable in the presence of GndHCl (66–70). These data indicate that CWD prions, regardless of the tissue of origin, are stable in the presence of denaturing agents.

We did observe differences in electrophoretic mobility of the obex and the lymph node samples treated with ≥ 2.5 M GndHCl that suggest more subtle differences in prion structural dynamics between prions present in these two tissues. All obex samples shift farther down the gel than lymph node samples when treated with 2.5 M or greater of GndHCl, indicating that neurogenic prions adopt a unique conformation at these chaotrope concentrations, allowing PK differential access to these prions compared to lymphogenic prions. Whether these apparent biochemical differences between neurogenic and lymphogenic prions in conformational stability and structural dynamics translate to biological significance remains to be determined. We also detected numerous high-molecular weight species staining for PrP, which may represent prion oligomers, in both obex and lymph node samples.

While we observed no differences between mean [GndHCl]_1/2_ values between prions isolated from lymph nodes compared to obex, we witnessed more variable conformational stability in lymph node-derived prion samples, as evidenced by large variances observed at each [GndHCl] (Figure 2). Moreover, we observed unequal variances between the brain and lymph node mean [GndHCl]_1/2_ value of one animal (10083) and a nearly significant difference in another (10074), indicating more variability in lymphogenic prion conformation. Finally, the denaturation curves appear different between the brain and lymph node samples, potentially revealing some biochemical differences between the two prion sources that aren’t entirely represented in the sample means (Figure 4).

We also found significant differences in glycoform ratios between the paired obex and lymph node-derived prions in all four animals tested. Glycoform ratio differences are an important indicator of different prion strains and another line of evidence that the prions present in the lymph node differ somewhat to those in the obex of the same animal. These differences could signify a biochemical strain difference between these animals; however, there is no systematic assessment of the glycosylation pattern of PrP^C^ in the lymph node or brain of white-tailed deer and it is possible there are different pools of differentially glycosylated PrP between these two tissues. In rodent models of scrapie, investigations of PrP^C^ glycosylation profiles in the brain suggest that glycosylation influences neuroinvasion, PrP^Sc^ deposition and neuropathological lesion profiles (71,72). Similar studies with mouse models of CWD or observational studies of infected deer could determine the biological relevance of our observed cerPrP^Sc^ glycosylation differences.

Samples from individual animals were then averaged together to assess any tissue differences between the obex and lymph node samples across animals. We found no statistical differences in conformational stability (Figure 3), but the variances between the samples differed significantly. We also observed significant differences in glycoform ratios between brain and lymph node-derived cerprP^Sc^ (Figure 5 and Tables 2 and 3). These results mirrored the results observed in individual deer, reinforcing that biochemical strain differences exist between obex and lymph node CWD prions.

Lastly, we assessed biochemical strain differences among PrP^Sc^ present in the same tissue across individuals to identify any differences among animals. We observed no differences in conformational stability among any of the obex or lymph node samples across individuals. These data suggest that while there are some indications of conformational differences among strains between tissues, there do not appear to be any significant conformational differences between PrP^Sc^ in tissues across individuals. When we compared PrP^Sc^ glycoform ratios from the same tissue type across individual deer, we observed limited differences in glycoform ratio in the lymph node samples (Figure 5 and Table 2). All significant differences occurred between sample 07416 and 10074 or 07399. Surprisingly, though, we observed many differences in obex PrP^Sc^ glycoform ratios among individual deer (figure 5 and Table 3).

Based on our hypothesis of greater strain diversity of lymphogenic prions, we propose a model in which diverse prion strains present in the lymphoid organs may traffic to the brain, where different PrP^C^ glycoforms present in different neuroanatomical regions select specific cerPrP^Sc^ isoforms and propagate specific of prion strains in those regions (Figure 6). This relatively more homogeneous group of prions produces predominant neurogenic prion strains that may be quite distinct among individual deer, like we observed in this study. The more variability that we see in lymph node prions may be due to a larger and more diverse population of prion strains in this extraneural site. We previously identified Complement proteins CD21/35 and Factor H as high affinity prion receptors in extraneural sites (73–75). Since the CNS express neither CD21/35 nor Factor H, but express far more PrP^C^ than the lymphoid system, PrP^C^ likely acts as the dominant prion receptor in the CNS and selects a more restricted set of prions, perhaps influenced by PrP^C^ glycosylation. Increased numbers and diversity of prion receptors outside the CNS may result in increased diversity of prion strains in extraneural tissues, as our biochemical data presented here indicates. Extraneural prions have been shown to have a wider species tropism that CNS prions(76). We propose that the increased prion diversity we measured biochemically in this work potentially increases their zoonotic potential as well.

**Figure 6.**
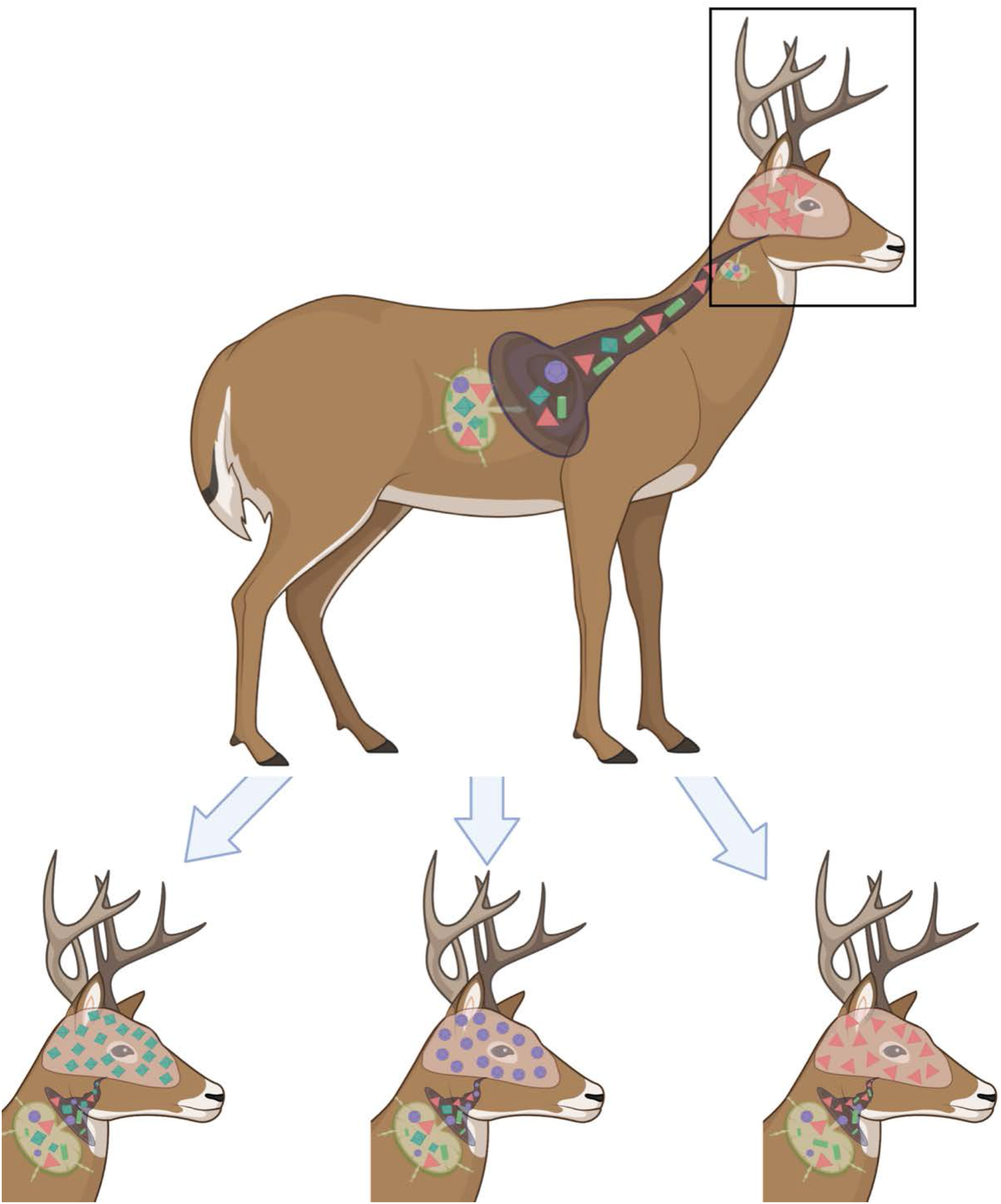
Proposed model of lymphoid prion replication influencing differential neural prion selection and propagation in individual cervids. Various lymphoid prion receptors, including Complement proteins CD21/35, Factor H, C3 and C1q, among others, select and replicate a more diverse pool of prions. PrP^C^, the predominant prion receptor in the CNS, selects a more restricted pool of prions, indicated by the narrowing “funnel” to the brain. These prions may differ significantly among individuals, resulting in cervids expressing different predominant strains, likely with variable zoonotic potential.

Taken together, the data presented here provide strong evidence for biochemical strain differences between obex and lymph node derived prions in these cervids. Mouse bioassays are underway to see if these biochemical differences translate to biological differences and greater species tropism with zoonotic potential. If so, these results suggest that research should focus on extraneural prions and prion strains as they differ from neurogenic prions, are more likely to be shed into the environment and expose other cervids and humans to prions with greater zoonotic potential. This proposed increased zoonotic potential of lymphogenic prions also has implications for best practices and policies regarding what tissues hunters and producers provide to agencies, and what diagnostic tests those agencies perform on those tissues to assess not just prion positivity, but also strain properties that may indicate zoonotic potential.

## References

1. Miller MW, Wild MA. Epidemiology of chronic wasting disease in captive white-tailed and mule deer. Journal of Wildlife Diseases. 2004;40(2):320–7.

2. Miller MW, Williams ES. Horizontal prion transmission in mule deer. Nature [Internet]. 2003 Sep [cited 2019 Sep 23];425(6953):35–6. Available from: http://www.nature.com/articles/425035a

3. Miller MW, Wild MA. Epidemiology of Chronic Wasting Disease in Captive White-Tailed and Mule Deer. Journal of Wildlife Diseases [Internet]. 2004;40(2):320–7. Available from: http://www.jwildlifedis.org/doi/10.7589/0090-3558-40.2.320

4. Miller MW, Conner MM. Epidemiology of chronic wasting disease in free-ranging mule deer: Spatial, temporal, and demographic influences on observed prevalence patterns. Journal of Wildlife Diseases [Internet]. 2005;41(2):275–90. Available from: http://internal-pdf//EpidemiologyOfChronicWastingDiseaseInFreeRangingMuleDeerSpatialT-2225030656/EpidemiologyOfChronicWastingDiseaseInFreeRangingMuleDeerSpatialTemporalAndDemographicInfluencesOnObservedPrevalencePatterns.pdf%5Cn%3CGotoISI%3E://WOS:00023

5. Miller MW, Hobbs NT, Tavener SJ. Dynamics of prion disease transmission in mule deer. Ecol Appl. 2006;16(6):2208–14.

6. Miller MW, Williams ES, McCarty CW, Spraker TR, Kreeger TJ, Larsen CT, et al. EPIZOOTIOLOGY OF CHRONIC WASTING DISEASE IN FREE-RANGING CERVIDS IN COLORADO AND WYOMING. Journal of Wildlife Diseases [Internet]. 2000 Oct 1 [cited 2019 Sep 16];36(4):676–90. Available from: http://www.jwildlifedis.org/doi/10.7589/0090-3558-36.4.676

7. Nalls A v., McNulty E, Hoover CE, Pulscher LA, Hoover EA, Mathiason CK. Infectious Prions in the Pregnancy Microenvironment of Chronic Wasting Disease-Infected Reeves’ Muntjac Deer. Journal of Virology. 2017;91(15):1–15.

8. Nalls A v., McNulty E, Powers J, Seelig DM, Hoover C, Haley NJ, et al. Mother to Offspring Transmission of Chronic Wasting Disease in Reeves’ Muntjac Deer. Baskakov I v., editor. PLoS ONE [Internet]. 2013 Aug 14 [cited 2019 Sep 16];8(8):e71844. Available from: https://dx.plos.org/10.1371/journal.pone.0071844

9. Selariu A, Powers JG, Nalls A, Brandhuber M, Mayfield A, Fullaway S, et al. In utero transmission and tissue distribution of chronic wasting disease-associated prions in free-ranging Rocky Mountain elk. Journal of General Virology. 2015;96(11):3444–55.

10. Henderson DM, Denkers ND, Hoover CE, Garbino N, Mathiason CK, Hoover EA. Longitudinal Detection of Prion Shedding in Saliva and Urine by Chronic Wasting Disease-Infected Deer by Real-Time Quaking-Induced Conversion. Journal of Virology. 2015;89(18):9338–47.

11. Mathiason CK, Powers JG, Dahmes SJ, Osborn DA, Miller K v., Warren RJ, et al. Infectious prions in the saliva and blood of deer with chronic wasting disease. Science. 2006;314(5796):133–6.

12. Mathiason CK, Cohen M, McManus L, Mitchell R, Aguilar-Calvo P, Consolacion G, et al. Silent Prions and Covert Prion Transmission. True-Krob HL, editor. PLOS Pathogens [Internet]. 2015 Dec 10 [cited 2016 Sep 1];11(12):e1005249. Available from: http://dx.plos.org/10.1371/journal.ppat.1005249

13. Angers RC, Seward TS, Napier D, Green M, Hoover E, Spraker T, et al. Chronic wasting disease prions in eik antler velvet. Emerging Infectious Diseases. 2009;15(5):696–703.

14. Sigurdson CJ, Williams ES, Miller MW, Spraker TR, O’Rourke KI, Hoover EA. Oral transmission and early lymphoid tropism of chronic wasting disease PrP(res) in mule deer fawns (Odocoileus hemionus). Journal of General Virology. 1999;80(10):2757–64.

15. Sigurdson CJ, Barillas-Mury C, Miller MW, Oesch B, M van Keulen LJ, M Langeveld JP, et al. PrP CWD lymphoid cell targets in early and advanced chronic wasting disease of mule deer [Internet]. Vol. 83, Journal of General Virology. 2019 [cited 2019 Apr 17]. Available from: www.microbiologyresearch.org

16. Nichols TA, Spraker TR, Rigg TD, Meyerett-Reid C, Hoover C, Michel B, et al. Intranasal Inoculation of White-Tailed Deer (Odocoileus virginianus) with Lyophilized Chronic Wasting Disease Prion Particulate Complexed to Montmorillonite Clay. Kincaid AE, editor. PLoS ONE [Internet]. 2013 May 9 [cited 2016 Sep 6];8(5):e62455. Available from: http://dx.plos.org/10.1371/journal.pone.0062455

17. Mabbott NA, MacPherson GG. Prions and their lethal journey to the brain. [Internet]. Vol. 4, Nature reviews. Microbiology. Nature Publishing Group; 2006 [cited 2021 Feb 26]. p. 201–11. Available from: www.nature.com/reviews/micro

18. Hoover CE, Davenport KA, Henderson DM, Denkers ND, Mathiason CK, Soto C, et al. Pathways of Prion Spread during Early Chronic Wasting Disease in Deer. Journal of Virology [Internet]. 2017 May 15 [cited 2021 Feb 26];91(10). Available from: https://pubmed.ncbi.nlm.nih.gov/28250130/

19. Bartz JC. Prion strain diversity. Cold Spring Harbor Perspectives in Medicine. 2016;6(12).

20. Morales R. Prion strains in mammals: Different conformations leading to disease Strain variation in prion diseases. 2017 [cited 2021 Feb 26]; Available from: https://doi.org/10.1371/journal.ppat.1006323

21. Morales R, Abid K, Soto C. The prion strain phenomenon: Molecular basis and unprecedented features. Vol. 1772, Biochimica et Biophysica Acta - Molecular Basis of Disease. Elsevier; 2007. p. 681–91.

22. Angers RC, Kang HE, Napier D, Browning S, Seward T, Mathiason C, et al. Prion strain mutation determined by prion protein conformational compatibility and primary structure. Science. 2010 May 28;328(5982):1154–8.

23. Herbst A, Velásquez CD, Triscott E, Aiken JM, McKenzie D. Chronic wasting disease prion strain emergence and host range expansion. Emerging Infectious Diseases. 2017;23(9):1598–600.

24. Monello RJ, Galloway NL, Powers JG, Madsen-Bouterse SA, Edwards WH, Wood ME, et al. Pathogen-mediated selection in free-ranging elk populations infected by chronic wasting disease. Proceedings of the National Academy of Sciences of the United States of America [Internet]. 2017 Nov 14 [cited 2019 Sep 16];114(46):12208–12. Available from: http://www.ncbi.nlm.nih.gov/pubmed/29087314

25. Benestad SL, Sarradin P, Thu B, Schönheit J, Tranulis MA, Bratberg B. Cases of scrapie with unusual features in Norway and designation of a new type, Nor98. Veterinary Record. 2003 Aug 16;153(7):202–8.

26. Chafin TK, Douglas MR, Martin BT, Zbinden ZD, Middaugh CR, Ballard JR, et al. Age structuring and spatial heterogeneity in prion protein gene (PRNP) polymorphism in white-tailed deer. https://doi.org/101080/1933689620201832947 [Internet]. 2020 Jan 1 [cited 2021 Jul 14];14(1):238–48. Available from: https://www.tandfonline.com/doi/abs/10.1080/19336896.2020.1832947

27. Béringue V, Herzog L, Jaumain E, Reine F, Sibille P, le Dur A, et al. Facilitated cross-species transmission of prions in extraneural tissue. Science [Internet]. 2012;335(6067):472–5. Available from: http://dx.doi.org/10.1126/science.1215659

28. Angers RC, Browning SR, Seward TS, Sigurdson CJ, Miller MW, Hoover EA, et al. Prions in skeletal muscles of deer with chronic wasting disease. Science. 2006 Feb 24;311(5764):1117.

29. Greenlee JJ, Nicholson EM, Smith JD, Kunkle RA, Hamir AN. Susceptibility of cattle to the agent of chronic wasting disease from elk after intracranial inoculation. Journal of Veterinary Diagnostic Investigation. 2012 Nov;24(6):1087–93.

30. Hamir AN, Kunkle RA, Cutlip RC, Miller JM, Williams ES, Richt JA. Transmission of chronic wasting disease of mule deer to Suffolk sheep following intracerebral inoculation. Journal of Veterinary Diagnostic Investigation. 2006;18(6):558–65.

31. Moore SJ, West Greenlee MH, Kondru N, Manne S, Smith JD, Kunkle RA, et al. Experimental Transmission of the Chronic Wasting Disease Agent to Swine after Oral or Intracranial Inoculation. Journal of Virology [Internet]. 2017 Oct 1 [cited 2021 Feb 26];91(19). Available from: https://doi.org/10.1128/JVI

32. Mathiason CK, Nalls A v., Seelig DM, Kraft SL, Carnes K, Anderson KR, et al. Susceptibility of Domestic Cats to Chronic Wasting Disease. Journal of Virology. 2013 Feb 15;87(4):1947–56.

33. Davenport KA, Christiansen JR, Bian J, Young M, Gallegos J, Kim S, et al. Comparative analysis of prions in nervous and lymphoid tissues of chronic wasting disease-infected cervids. Journal of General Virology [Internet]. 2018 May 1 [cited 2021 Apr 17];99(5):753–8. Available from: https://www.microbiologyresearch.org/content/journal/jgv/10.1099/jgv.0.001053

34. Michel B, Ferguson A, Johnson T, Bender H, Meyerett-Reid C, Wyckoff AC, et al. Complement protein C3 exacerbates prion disease in a mouse model of chronic wasting disease. International Immunology [Internet]. 2013 Dec 1 [cited 2016 Sep 1];25(12):697– 702. Available from: http://www.intimm.oxfordjournals.org/cgi/doi/10.1093/intimm/dxt034

35. Zabel MD, Heikenwalder M, Prinz M, Arrighi I, Schwarz P, Kranich J, et al. Stromal complement receptor CD21/35 facilitates lymphoid prion colonization and pathogenesis. Journal of immunology (Baltimore, Md : 1950) [Internet]. 2007 Nov 1 [cited 2019 Apr 21];179(9):6144–52. Available from: http://www.ncbi.nlm.nih.gov/pubmed/17947689

36. Michel B, Ferguson A, Johnson T, Bender H, Meyerett-Reid C, Pulford B, et al. Genetic Depletion of Complement Receptors CD21/35 Prevents Terminal Prion Disease in a Mouse Model of Chronic Wasting Disease. The Journal of Immunology [Internet]. 2012 Nov 1 [cited 2016 Sep 1];189(9):4520–7. Available from: http://www.jimmunol.org/cgi/doi/10.4049/jimmunol.1201579

37. Kane SJ, Farley TK, Gordon EO, Estep J, Bender HR, Moreno JA, et al. Complement Regulatory Protein Factor H Is a Soluble Prion Receptor That Potentiates Peripheral Prion Pathogenesis. The Journal of Immunology [Internet]. 2017;ji1701100. Available from: http://www.jimmunol.org/lookup/doi/10.4049/jimmunol.1701100

38. Kane SJ, Swanson E, Gordon EO, Rocha S, Bender HR, Donius LR, et al. Relative Impact of Complement Receptors CD21/35 (Cr2/1) on Scrapie Pathogenesis in Mice. mSphere [Internet]. 2017 Nov 22 [cited 2021 Feb 26];2(6). Available from: https://pubmed.ncbi.nlm.nih.gov/29202042/

39. Mabbott NA, Bruce ME. Complement component C5 is not involved in scrapie pathogenesis. Immunobiology [Internet]. 2004 Nov 9 [cited 2019 Apr 17];209(7):545–9. Available from: https://www.sciencedirect.com/science/article/pii/S0171298504000695?via%3Dihub

40. Bradford BM, Crocker PR, Mabbott NA. Peripheral prion disease pathogenesis is unaltered in the absence of sialoadhesin (Siglec-1/CD169). Immunology [Internet]. 2014 Sep 1 [cited 2019 Apr 17];143(1):120–9. Available from: http://doi.wiley.com/10.1111/imm.12294

41. Barria MA, Telling GC, Gambetti P, Mastrianni JA, Soto C. Generation of a new form of human PrPScin vitro by interspecies transmission from cervid prions. Journal of Biological Chemistry. 2011;286(9):7490–5.

42. Barria MA, Balachandran A, Morita M, Kitamoto T, Barron R, Manson J, et al. Molecular barriers to zoonotic transmission of prions. Emerging Infectious Diseases. 2014;20(1):88– 97.

43. Crowell J, Hughson A, Caughey B, Bessen RA. Host Determinants of Prion Strain Diversity Independent of Prion Protein Genotype. Beemon KL, editor. Journal of Virology [Internet]. 2015 Oct 15 [cited 2016 Sep 1];89(20):10427–41. Available from: http://jvi.asm.org/lookup/doi/10.1128/JVI.01586-15

44. le Dur A, Laï TL, Stinnakre MG, Laisné A, Chenais N, Rakotobe S, et al. Divergent prion strain evolution driven by PrP C expression level in transgenic mice. Nature Communications [Internet]. 2017 Jan 23 [cited 2021 Mar 31];8(1):1–11. Available from: www.nature.com/naturecommunications

45. Bessen RA, Marsh RF. Biochemical and Physical Properties of the Prion Protein from Two Strains of the Transmissible Mink Encephalopathy Agent [Internet]. JOURNAL OF VIROLOGY. 1992. Available from: http://jvi.asm.org/

46. Bessen RA, Marsh RF. Distinct PrP properties suggest the molecular basis of strain variation in transmissible mink encephalopathy. Journal of Virology [Internet]. 1994 [cited 2021 Apr 14];68(12):7859–68. Available from: https://pubmed.ncbi.nlm.nih.gov/7966576/

47. Hill AF, Desbruslais M, Joiner S, Sidle KCL, Gowland I, Collinge J, et al. The same prion strain causes vCJD and BSE. Nature [Internet]. 1997 Oct 2 [cited 2018 Jul 9];389(6650):448–50. Available from: http://www.nature.com/articles/38925

48. Velásquez CD, Chae K, Tracy H, Chiye K, Allen H, Judd A, et al. Chronic wasting disease (CWD) prion strains evolve via adaptive diversification of conformers in hosts expressing prion protein polymorphisms. J Biol Chem. 2020;295(15):4985–5001.

49. Parchi P, de Boni L, Saverioni D, Cohen ML, Ferrer I, Gambetti P, et al. Consensus classification of human prion disease histotypes allows reliable identification of molecular subtypes: An inter-rater study among surveillance centres in Europe and USA. Acta Neuropathologica [Internet]. 2012 Oct 30 [cited 2021 Apr 15];124(4):517–29. Available from: https://link.springer.com/article/10.1007/s00401-012-1002-8

50. Edmunds DR, Kauffman MJ, Schumaker BA, Lindzey FG, Cook WE, Kreeger TJ, et al. Chronic wasting disease drives population decline of white-tailed deer. PLoS ONE. 2016;11(8):1–19.

51. DeVivo MT, Edmunds DR, Kauffman MJ, Schumaker BA, Binfet J, Kreeger TJ, et al. Endemic chronic wasting disease causes mule deer population decline in Wyoming. Zabel MD, editor. PLOS ONE [Internet]. 2017 Oct 19 [cited 2019 Sep 16];12(10):e0186512. Available from: https://dx.plos.org/10.1371/journal.pone.0186512

52. Manjerovic MB, Green ML, Mateus-Pinilla N, Novakofski J. The importance of localized culling in stabilizing chronic wasting disease prevalence in white-tailed deer populations. Preventive Veterinary Medicine [Internet]. 2014;113(1):139–45. Available from: http://dx.doi.org/10.1016/j.prevetmed.2013.09.011

53. Almberg ES, Cross PC, Johnson CJ, Heisey DM, Richards BJ. Modeling routes of chronic wasting disease transmission: Environmental prion persistence promotes deer population decline and extinction. PLoS ONE. 2011;6(5).

54. Uehlinger FD, Johnston AC, Bollinger TK, Waldner CL. Systematic review of management strategies to control chronic wasting disease in wild deer populations in North America. BMC Veterinary Research [Internet]. 2016;12(1):173. Available from: http://bmcvetres.biomedcentral.com/articles/10.1186/s12917-016-0804-7

55. Bendheim PE, Brown HR, Rudelli RD, Scala LJ, Goller NL, Wen GY, et al. Nearly ubiquitous tissue distribution of the scrapie agent precursor protein. Neurology [Internet]. 1992 Jan 1 [cited 2021 Mar 3];42(1):149–56. Available from: https://n.neurology.org/content/42/1/149

56. Ford MJ, Burton LJ, Morris RJ, Hall SM. Selective expression of prion protein in peripheral tissues of the adult mouse. Neuroscience. 2002 Aug 2;113(1):177–92.

57. Davenport KA, Hoover CE, Bian J, Telling GC, Mathiason CK, Hoover EA. PrP C expression and prion seeding activity in the alimentary tract and lymphoid tissue of deer. 2017; Available from: https://doi.org/10.1371/journal.pone.0183927

58. Miele G, Alejo Blanco AR, Baybutt H, Horvat S, Manson J, Clinton M. Embryonic activation and developmental expression of the murine prion protein gene. Gene Expression [Internet]. 2003 [cited 2021 Apr 25];11(1):1–12. Available from: https://pubmed.ncbi.nlm.nih.gov/12691521/

59. Fournier JG, Escaig-Haye F, Billette De Villemeur T, Robain O, Lasmézas CI, Deslys JP, et al. Distribution and submicroscopic immunogold localization of cellular prion protein (PrPc) in extracerebral tissues. Cell and Tissue Research [Internet]. 1998 [cited 2021 Apr 25];292(1):77–84. Available from: https://pubmed.ncbi.nlm.nih.gov/9506914/

60. Horiuchi M, Yamazaki N, Ikeda T, Ishiguro N, Shinagawa M. A cellular form of prion protein (PrP(C)) exists in many non-neuronal tissues of sheep. Journal of General Virology [Internet]. 1995 Oct 1 [cited 2021 Apr 25];76(10):2583–7. Available from: https://www.microbiologyresearch.org/content/journal/jgv/10.1099/0022-1317-76-10-2583

61. Prusiner SB, Scott MR, DeArmond SJ, Cohen FE. Prion protein biology [Internet]. Vol. 93, Cell. Elsevier B.V.; 1998 [cited 2021 Apr 25]. p. 337–48. Available from: http://www.cell.com/article/S0092867400811630/fulltext

62. Liu T, Li R, Wong B-S, Liu D, Pan T, Petersen RB, et al. Normal Cellular Prior Protein Is Preferentially Expressed on Subpopulations of Murine Hemopoietic Cells. The Journal of Immunology [Internet]. 2001 Mar 15 [cited 2021 Apr 25];166(6):3733–42. Available from: https://pubmed.ncbi.nlm.nih.gov/11238614/

63. Oesch B, Westaway D, Wälchli M, McKinley MP, Kent SBH, Aebersold R, et al. A cellular gene encodes scrapie PrP 27-30 protein. Cell [Internet]. 1985 [cited 2021 Apr 14];40(4):735–46. Available from: https://pubmed.ncbi.nlm.nih.gov/2859120/

64. Fox KA, Jewell JE, Williams ES, Miller MW. Patterns of PrPCWD accumulation during the course of chronic wasting disease infection in orally inoculated mule deer (Odocoileus hemionus). Journal of General Virology [Internet]. 2006 Nov 1 [cited 2019 Sep 16];87(11):3451–61. Available from: https://www.microbiologyresearch.org/content/journal/jgv/10.1099/vir.0.81999-0

65. Selariu A, Powers JG, Nalls A, Brandhuber M, Mayfield A, Fullaway S, et al. In utero transmission and tissue distribution of chronic wasting disease-associated prions in free-ranging Rocky Mountain elk. Journal of General Virology. 2015;96(11):3444–55.

66. Meyerett C, Michel B, Pulford B, Spraker TR, Nichols TA, Johnson T, et al. In vitro strain adaptation of CWD prions by serial protein misfolding cyclic amplification. Virology. 2008;382(2).

67. Meyerett-Reid C, Wyckoff AC, Spraker T, Pulford B, Bender H, Zabel MD. De novo generation of a unique cervid prion strain using protein misfolding cyclic amplification. mSphere. 2017;2(1).

68. Xie Z, Rourke KIO, Dong Z, Jenny AL, Langenberg JA, Belay ED, et al. Chronic Wasting Disease of Elk and Deer and Creutzfeldt-Jakob Disease COMPARATIVE ANALYSIS OF THE SCRAPIE PRION PROTEIN *. 2006;281(7):4199–206.

69. Green KM, Browning SR, Seward TS, Jewell JE, Ross DL, Green MA, et al. The elk PRNP codon 132 polymorphism controls cervid and scrapie prion propagation. Journal of General Virology. 2008;89(2):598–608.

70. Scott MR, Peretz D, Nguyen H-OB, DeArmond SJ, Prusiner SB. Transmission Barriers for Bovine, Ovine, and Human Prions in Transgenic Mice. Journal of Virology. 2005;79(9):5259–71.

71. Cancellotti E, Bradford BM, Tuzi NL, Hickey RD, Brown D, Brown KL, et al. Glycosylation of PrPC Determines Timing of Neuroinvasion and Targeting in the Brain following Transmissible Spongiform Encephalopathy Infection by a Peripheral Route. Journal of Virology. 2010;84(7):3464–75.

72. DeArmond SJ, Qiu Y, Sanchez H, Spilman PR, Ninchak-Casey A, Alonso D, et al. PrPc glycoform heterogeneity as a function of brain region: implications for selective targeting of neurons by prion strains. J Neuropathol Exp Neurol. 1999;58(9):1000–9.

73. Michel B, Ferguson A, Johnson T, Bender H, Meyerett-Reid C, Pulford B, et al. Genetic depletion of complement receptors CD21/35 prevents terminal prion disease in a mouse model of chronic wasting disease. Journal of Immunology. 2012;189(9).

74. Kane SJ, Farley TK, Gordon EO, Estep J, Bender HR, Moreno JA, et al. Complement Regulatory Protein Factor H Is a Soluble Prion Receptor That Potentiates Peripheral Prion Pathogenesis. The Journal of Immunology. 2017;199(11):3821–7.

75. Kane SJ, Swanson E, Gordon EO, Rocha S, Bender HR, Donius LR, et al. Relative impact of complement receptors CD21/35 (Cr2/1) on scrapie pathogenesis in mice. mSphere. 2017;2(6).

76. Béringue V, Herzog L, Jaumain E, Reine F, Sibille P, Le Dur A, et al. Facilitated cross-species transmission of prions in extraneural tissue. Science. 2012;335(6067):472–5.

